# NETosis Induced by Serum of Patients with COVID-19 is Reduced with Reparixin or Antibodies Against DEK and IL-8

**DOI:** 10.1101/2023.03.30.534873

**Authors:** İrfan Baki Kılıç, Açelya Yaşar, İrem Yalım Camcı, Türkan Güzel, Ayşegül Karahasan, Tamer Yağcı, Naci Çine, Ayten Kandilci

## Abstract

DEK locates in the nucleus of the cells or the cytoplasmic granules of neutrophils and plays different roles in cellular processes including NETosis, a suicide mechanism of neutrophils. Here we showed that the interaction of rDEK with CXCR2 leads to NETosis, which could be reduced by the CXCR1/CXCR2 inhibitor reparixin. We found that IL-8, IL-6, IL1-β, MPO, and CitH3 were increased whereas DEK was decreased in the serum of COVID-19 patients. Interestingly, reparixin or anti-DEK antibody reduced the NETosis induced by the serums of patients, suggesting that initial cytokine stimulation may further induce the release of DEK. Our results support the use of reparixin as a potential therapeutic strategy in COVID-19 and suggest that DEK-CXCR2 interaction plays a role in NETosis.

## INTRODUCTION

NETosis is characterized by extracellular web-like structures called neutrophil extracellular traps (NETs) that are generated with cell-extruded DNA bound to histones and granule proteins [1]. Microbial or viral components and pro-inflammatory cytokines may trigger NETosis. Although it is an important immune defense mechanism against pathogens, uncontrolled NETosis contributes to severe tissue damage leading to acute lung injury in influenza or SARS-CoV-2 infections. Neutrophils infected with SARS-CoV-2, and cytokine storm caused by this infection induce NETosis in COVID-19, which may be fatal in 1-3% of patients who were not vaccinated [2–7]. Among the cytokines, IL-8 is a potent inducer of NETosis, and patients with COVID-19 exhibit a high level of IL-8 in their serum and lung tissues [3, 8, 9].

Ubiquitously expressed oncoprotein DEK shares an ELR motive with IL-8, and it acts as a chemokine for neutrophils, CD8+ T-cells, and NK cells when secreted [10, 11]. Accumulating data point to DEK as a player in inflammation. A wide range of post-translational modifications influences multiple roles of DEK, mainly via decreasing its DNA binding capacity and allowing its secretion [12]. DEK is secreted passively from T-lymphocytes that undergo apoptosis and actively from immune cells and epithelial cells after IL-8 stimulation [10–13]. Secreted DEK may interact with heparin sulfate peptidoglycan receptors leading to entrance into the cell [10, 14], or it may interact with the CXCR2 receptor and via Gαi signaling suppresses hematopoietic progenitor cell proliferation [15]. Patients with JIA carry autoantibodies against DEK, and neutrophils from the Dek-knockout JIA mouse model have reduced capacity to form NETs after PMA or LPS stimulation; however, they retrieve to NETosis when co-stimulated with rDEK and PMA (or LPS) in vitro. Importantly, DEK-targeting aptamers reduce NETosis of the neutrophils in the JIA-mouse model [16]. In addition, human bronchial epithelial cells release DEK upon IL-8 stimulation, and supernatants of these cells, or rDEK alone, induce NETosis of human neutrophils in vitro, which is suppressed by overexpression of miR-181b-5p that directly targets and inhibits DEK mRNA expression [13].

Even though the current data suggest that DEK contributes to NETosis via regulating the chromatin architecture [16], the role of DEK-CXCR2 interaction in NETs formation is not elucidated yet. Here we aimed to determine whether the interaction of rDEK with the CXCR2 receptor affects NETosis in vitro. Given that NETosis is an important contributor to COVID-19 pathogenesis, we also analyzed and compared the amount of DEK and the selected cytokines elevated in the circulation of patients with COVID-19. Furthermore, we tested the inhibitory effect of reparixin, anti-DEK, and anti-IL-8 antibodies on the NETosis induced by COVID-19 patients’ serums. Our results indicate that DEK-CXCR2 interaction contributes to NETosis and support the therapeutic use of reparixin in COVID-19-related NETosis.

## METHOD

### Human Samples

Serum samples of COVID-19-patients (n=67; obtained from Marmara University Pendik Training and Research Hospital) or healthy volunteers (n=47; from Kocaeli University, Faculty of Medicine) were collected, aliquoted and stored at −20□. Patients’ samples were grouped as acute phase (n= 23; average age= 51.2 ±21 (mean ± SD); 12 males/11 females; serums were obtained 24-48 hours after RT-PCR positivity) and convalescent phase samples (n=44, average age= 53.5 ±12.2 (mean ± SD); 22 males/22 females). All patients were tested positive for SARS-CoV-2 at the hospital. Briefly, viral RNA was extracted from nasopharyngeal swab samples by using Bio-speedy® viral nucleic acid buffer (Bioexen LTD, Turkey) and RT-PCR was performed with Bio-speedy® COVID-19 qPCR detection kit, Version 2 (Bioexen LTD) using primers and probes targeting the RNA-dependent RNA polymerase (RdRp) gene fragment in a LightCycler® 96 System (Roche, Switzerland). Healthy donors’ samples were confirmed negative by using a COVID-19 IgG/IgM lateral flow antibody test (UNSCIENCE, UnCov-40). This study was approved by the ethics committee of Kocaeli University Medical Faculty (GOKAEK-2020/11.14 and GOKAEK-2021/15.13). Study was performed in accordance with the Declaration of Helsinki.

### ELISA Test

The level of the DEK, IL-6 (Both from Cusabio), IL-1β, MPO, IL8 (All three from Abcam) and Cit-H3 (Cayman) in the serum samples were detected by using ELISA kits and applying manufacturers’ protocol. Serums were diluted 1:5 for DEK, IL-1β, IL6, Cit-H3, and 1:8 for IL-8 and MPO before using.

### Neutrophil Isolation and NETosis Assays

In all experiments, neutrophils of healthy volunteers were used. Neutrophils were isolated by following the protocol published by Brinkmann et al., 2010 [17]. 2×10^5^ neutrophils/well were seeded into a poly-l-lysin (100 µg/ml, Sigma Aldrich, P5899) coated 8-well IBIDI slide (IBIDI, 80827) or a 96-well plate in culture medium (RPMI 1640 containing %1 human serum). After addition of recombinant human proteins (DEK, MyBioSource; MBS968198; IL-8, R&D 208-IL8/CF) or in some experiments patients’ serum (5% of the total volume of the culture medium [7]), cells were treated for an additional 1hr (for recombinant proteins) or for 2 hrs (for patients’ serum). When necessary, the cells were pre-treated with the indicated antibodies or reparixin (Med Chem, HY-15251) for indicated times in the legends. Then the cell-impermeable Sytox (Thermo Fisher, S7020) (at final concentration of 500 nM) was added to the cells for additional 15 min. Cells were fixed with 4% PFA (Sigma Aldrich, P6148) in the presence of Sytox, washed with PBS and the images were captured by using confocal microscope (Zeiss, LSM880) or ZOE Fluorescence Cell Imager (Biorad). Five different areas per well were imaged in the IBIDI or 96-well plates. Anti-DEK antibodies obtained from BD Biosciences (610948) and Bethyl Laboratories (A301-335A) whereas anti-IL-8 was from R&D (MAB208). All chemicals were first tested for the optimal dose (data not shown) and the neutrophils were treated with each corresponding vehicle (or with %5 healthy serum in the patients’ serum induced NETosis assays) as a negative control in all experiments.

For the quantification of NETosis, Image J (National Institutes of Health) and DANA I (DNA Area and NETosis Analysis) software was used [18]. In this analysis, NETs were quantified by comparing and counting the cell areas to the control cells with a normal morphology using cut-off 2 [18].

### Quantification of NET-associated MPO activity

MPO activity was detected by following the protocol published by Zuo et al., 2020 [7]. 1×10^5^ neutrophils were seeded into the 96-well plate and stimulated for NETosis as indicated above. After stimulation, medium was discarded. Cells were treated 5U/ml Micrococcal Nuclease (Thermo Fisher, EN0181) in 1XPBS for 10 minutes at 37□ in an incubator with 5% CO_2_. Reaction was stopped by adding 10 mM EDTA (final concentration 0.5 mM) (Invitrogen, AM9261) and the PBS containing NETs was transferred into a V-shaped 96-well plate. After centrifugation at 400g for 5 minutes, supernatant transferred into a new F-bottom 96-well plate and the same volume of TMB substrate (Thermo Fisher, 34021) was added to each well. The plate was incubated at room temperature for 10 min and the reaction was stopped by adding the same volume of 2N sulfuric acid. Absorbance was measured at 450nm by using a MultiSkan Fc (Thermo Fisher).

### Immunofluorescence Staining

Neutrophils were seeded into the 8 well IBIDI µ-slides as 2×10^5^ cells per well. NETosis was stimulated as described above with a final 15 min incubation in the presence of Sytox. After adding an equal amount of 8% PFA (final concentration 4%) [17] and fixing the cells for overnight at 4^0^C, cells were washed with PBS, permeabilized with 0,5% Triton-X-100 (Sigma Aldrich, T8787) for 1 min and blocked with 2% BSA (Sigma Aldrich, A2153) prepared in PBS for 20 min. Subsequently, cells were stained with a 1:50 diluted rabbit anti human DEK antibody (Bethyl Laboratories, A301-335A) for 1 hr. Then the cells were washed and incubated for additional 1 hr with a 1:500 diluted secondary anti rabbit Alexa Fluor 647 antibody (Cell Signalling Technologies, 4414). Then, the same cells were washed, fixed again with 4% PFA, blocked with 2% BSA and stained with a 1:50 diluted mouse anti human NE antibody (mouse, Thermo Fisher, MA1 40220) for overnight at 4^0^C. After staining for 1 hr with a 1:500 diluted anti mouse Alexa Fluor-555 secondary antibody (Cell Signaling Technologies, 4409), cells were directly imaged by using confocal microscopy.

### Statistical Analysis

All statistical analyses were performed by using GraphPad Prism (Version 8.0, GraphPad Software Inc). Significance was evaluated by Mann-Whitney test, Pearson and unpaired t-tests (ns, non-significant; ∗p < 0.05, ∗∗p < 0.01, ∗∗∗p < 0.001,).

## RESULTS

### Recombinant DEK induces NETosis via CXCR2 interaction

Including IL-8, all chemokines that bind CXCR1/CXCR2 receptors induce NETosis, and reparixin (an allosteric inhibitor of CXCR1/CXCR2) reduces NETs formation induced by these chemokines [8, 19]. To determine whether NETosis induced via rDEK also depends on CXCR2 interaction, we induced the primary neutrophils with either rDEK or rIL-8 as a control and then analyzed the NETs associated markers including the morphological changes of the cells, the presence of extracellular neutrophil elastase (NE), and the activity of NETs-associated myeloperoxidase (MPO). We found that both rDEK and rIL-8 (Figure 1A, 1C, 1D) stimulate NETs formation and pre-incubation of the cells with antibodies against DEK or IL-8 suppress this cellular process (Fig 1B, 1C, 1D). Interestingly, pre-treatment with reparixin also reduced DEK-induced NETosis indicating that DEK-CXCR2 interaction plays a role in NETs formation (Fig 1B, 1C, 1D).

**Figure 1:**
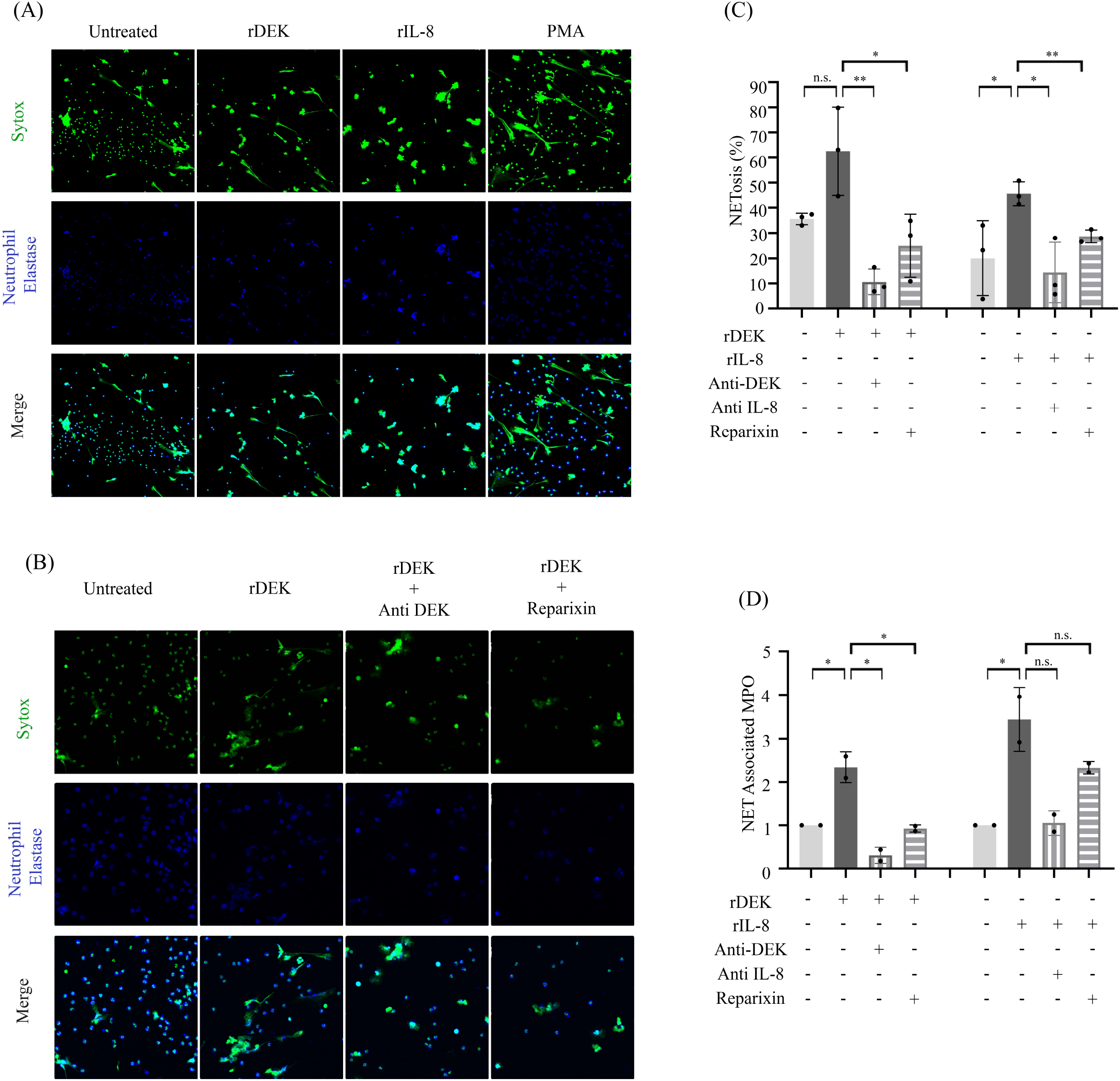
Reparixin reduces DEK-induced NETosis. Primary neutrophils were induced with rDEK (5 µg/ml (A) or 3.5 µg/ml (B, C, D) or rIL-8 (100ng/ml) for 1 hr; Sytox was added and the cells were incubated for an additional 15 min. For the confocal microscopy images (200X) in (A) and (B), cells were fixed and further stained with anti-NE antibody (blue). In a parallel experimental group, cells were first pretreated with anti-DEK (5 µg/ml in B; 2.5 µg/ml in C and D), anti-IL-8 (2.5 µg/ml), or reparixin (400 nM in B; 200 nM in C and D) for 15 min. (C) Confocal images captured in 5 different areas were analyzed using ImageJ and DANAI software and the percentage of cells showing NETs morphology was shown in the graph. Black circles in the graphs indicate the results of three independent assays. (D) Cells were treated similarly with recombinant proteins or chemical agents and NETs-associated MPO activity were measured. Black circles in the graphs indicate the results of two independent assays. Error bars: ±SD; NS: non-significant; *P < 0.05; **P < 0.01; ***P < 0.001 (Parametric unpaired t-test).

### The Level of DEK is lower in the serum of patients with COVID-19

Next, we analyzed whether the level of DEK in the circulation of patients was affected along with cytokines (IL-8, IL-6, IL-1β) and NETosis markers (MPO, CitH3) by using ELISA. We found that the DEK was significantly lower whereas all other analyzed markers were significantly higher in the serums of patients (n=67) compared to the healthy controls (n=38) (Figure 2A). When we grouped patients, we observed that the level of DEK was lower at the early stage (24 to 48 hours after the positive PCR-test result; called acute phase (AF) (n=23)) when compared to the late stage of the disease (21 days after the initial positive PCR-test; convalescent phase (CF)(n=44)) (Figure 2B). Contrary to the DEK, the level of all other markers was higher at CF compared to the AF of the disease, as reported before (Figure 2C-2G) [20]. Pearson correlation analysis indicated a significant negative correlation only between the level of DEK and IL-8 in healthy individuals (Figure 2H) and between the level of DEK and MPO in the AF of COVID-19 (Figure 2I). Altogether, these results suggested that expression, translation and/or secretion of DEK might be affected upon exposure to SARS-CoV-2.

**Figure 2:**
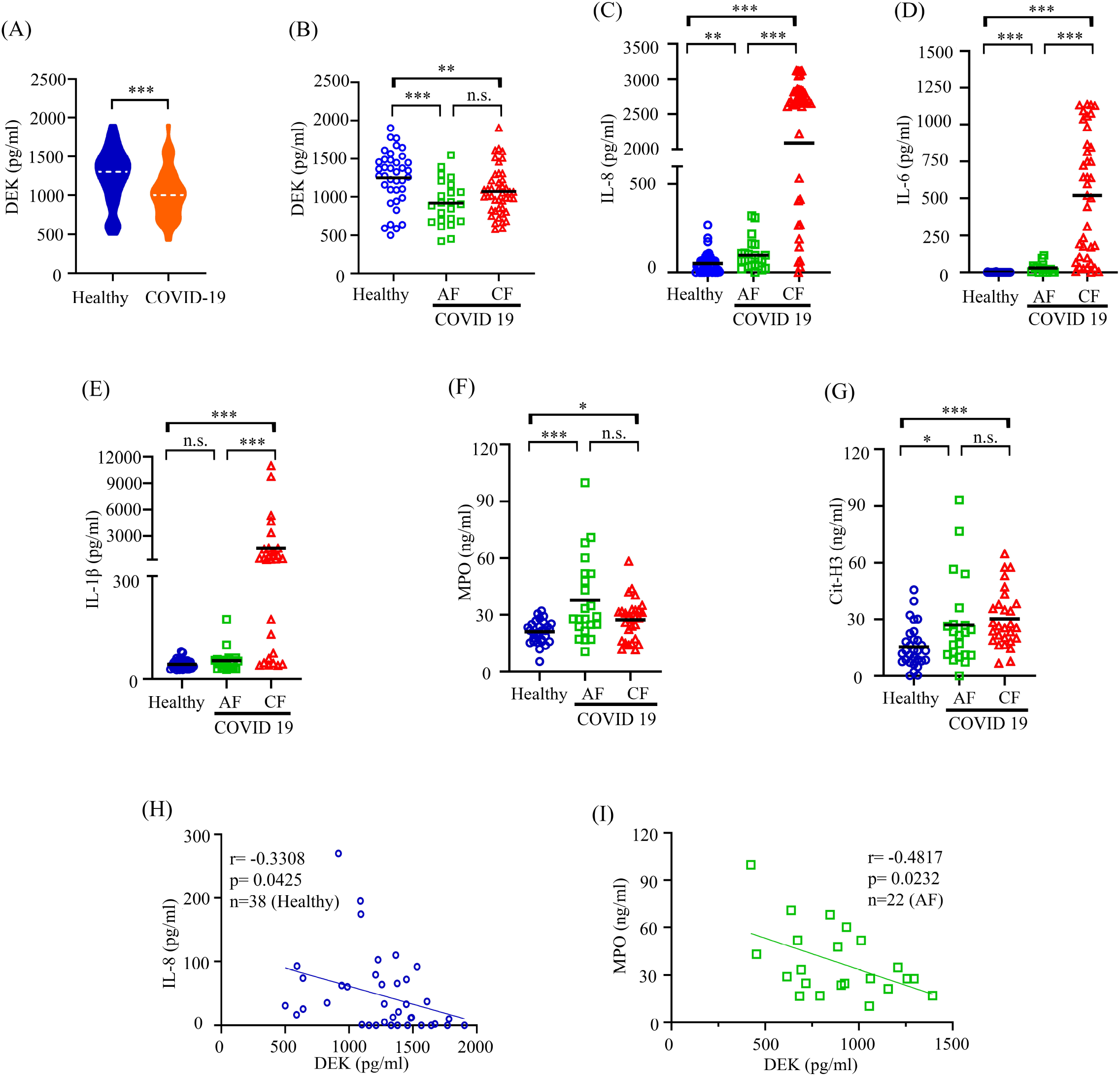
Patients with COVID-19 exhibit a lower level of DEK in the serum. (A-G) Level of DEK, IL-8 (healthy n=47; AF n= 23; CF n=44), IL-6 (healthy n=47; AF n= 16; CF n=39), IL-1β, MPO and CitH3 (healthy n=30; AF n= 22; CF n=28) were detected by using ELISA kits. (H-I) Graphs show Pearson correlation between the level of DEK and IL-8 in healthy individuals (H) and in patients with COVID-19 (at the AF of the disease). Lines indicate mean values. n.s.: non-significant; *P < 0.05; **P < 0.01; ***P < 0.001 (Mann-Whitney test).

### Reparixin or anti-DEK and anti-IL-8 antibodies suppress NETosis stimulated with the serum of patients

Given that neutrophils treated with serum samples of patients with COVID-19 develop NETosis [7], we further examined the inhibitory effect of reparixin, anti-DEK, and anti-IL-8 antibodies on NETs formation induced by the patients’ serums. Consistent with the literature [7], we showed that neutrophils treated with the patients’ serums form NETs (Figures 3A and 3B). Immunofluorescence staining indicated that both DEK and NE locate in the granules of neutrophils (Figures 3A and 3B) and interestingly we didn’t observe DEK in the nucleus. In the cells stimulated with a patient’s serum, cell-extruded DNA showed staining with both DNA-binding dye Sytox and NE, whereas the DEK was contained mainly in the granule-like structures and barely co-located with Sytox or NE (Figure 3A and 3B). These results suggest that DEK mostly locates in the granules but not in the nucleus of neutrophils, given that this commercial polyclonal anti-DEK antibody labels also nuclear DEK in other cell types (Bethyl Laboratories A301-335A;https://www.fishersci.com/shop/products/dek-ihc-polyclonal-bethyl-laboratories-2/501571807). Further analysis of available serum samples of patients (n=15) similarly indicated that COVID-19-patients’ serum induces NETosis and all three agents (reparixin, anti-DEK, and anti-IL-8) significantly reduce the NETs formation, as judged by both morphological analysis of the cells and NETs-associated MPO activity (Figure 4A). When we grouped the patient samples based on the ELISA values of DEK and IL-8 (DEK high/IL-8 high; DEK high/IL-8 low; DEK low/IL-8 high) (Table S1), we noticed that anti-DEK or anti-IL-8 antibodies more effectively reduced the NETosis in the groups where their level was higher, whereas reparixin exhibited a better inhibition in all three groups (Figure 4B-4D).

**Figure 3:**
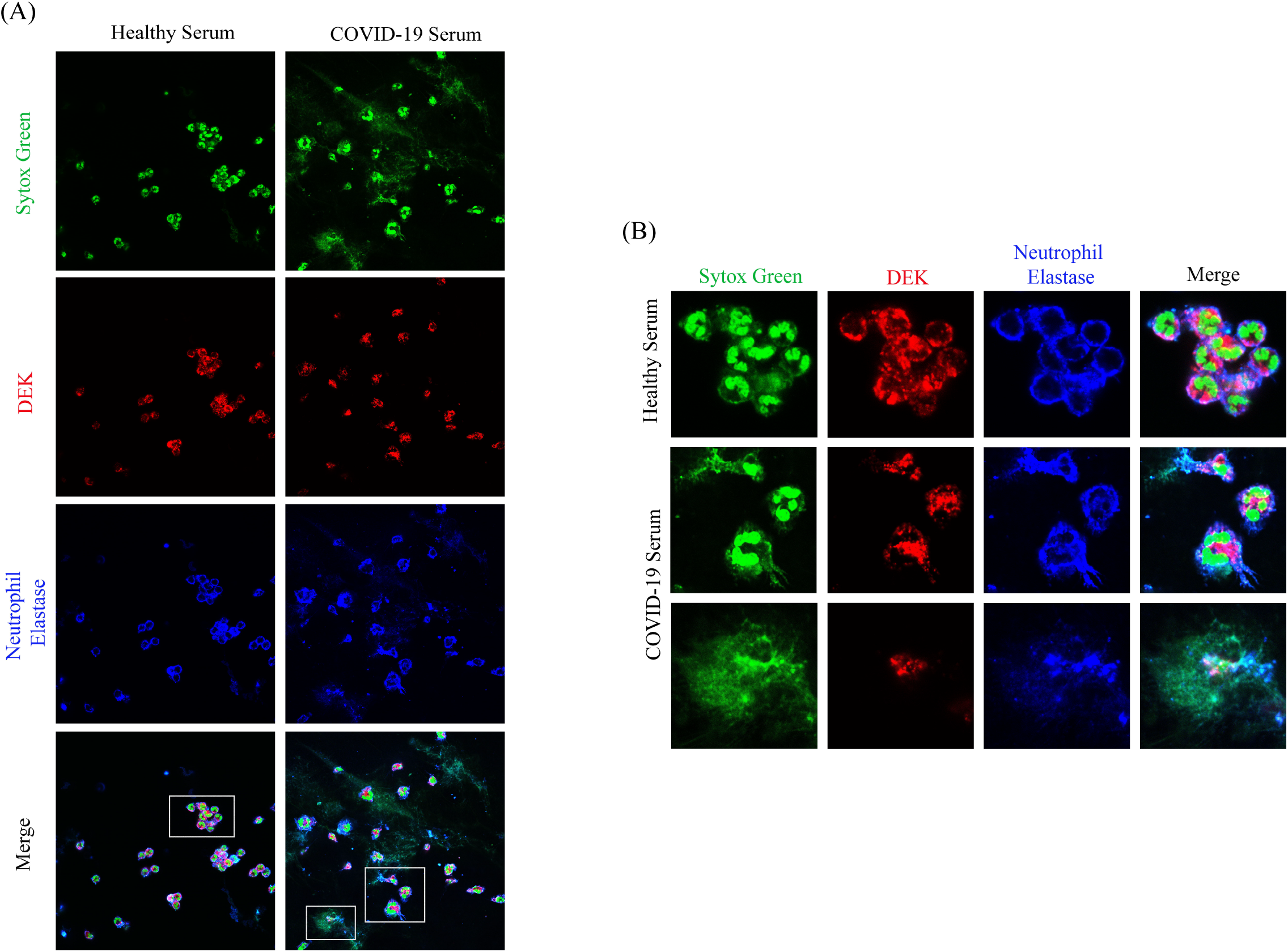
Immunofluorescence staining of neutrophils treated with healthy or COVID-19 serum samples. (A-B) Primary neutrophils were treated with 5% serum (healthy or COVID-19 samples) for two hrs then the cells were treated for an additional 15 min in the presence of the Sytox (green). Fixed cells were double-stained with rabbit anti-DEK (Bethyl Laboratories) (red) and mouse anti-NE (blue) antibodies. Images were taken by using a confocal microscope (630X, with oil). (B) Magnified images of the areas framed in (A) were shown.

**Figure 4:**
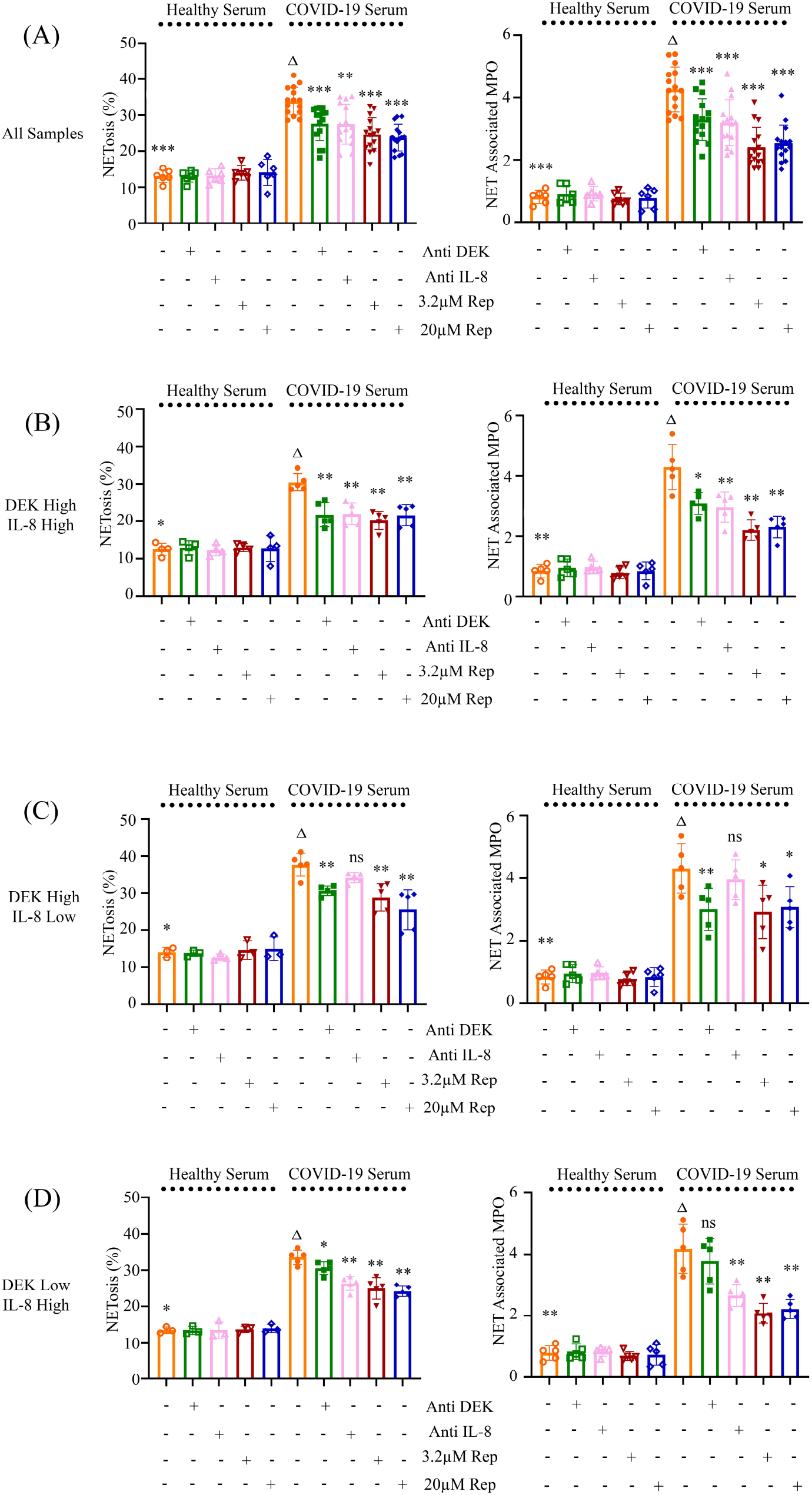
Reparixin or antibodies against DEK and IL-8 reduce NETosis induced by COVID-19-patients’ serum. (A-D) Neutrophils were treated with control (5% healthy serum) or COVID-19 serum (5%) samples for two hrs then the cells were treated for an additional 15 min in the presence of the Sytox. In an additional setting, cells were pretreated for two hrs with either reparixin (Rep), anti-DEK (Bethyl Laboratories), or anti-IL-8 antibodies before the addition of the 5% serum. Percent-NETosis analysis of the images taken from five different areas was performed by using ImageJ and DANAI software. For MPO analysis, cells were cultured similarly without the addition of sytox as described in the text. Graphs indicate the percentage of NETosis (left panels) and values of NETs associated MPO (right panels) in all of the analyzed samples [COVID-19 serum (n=15); healthy serum (n=4; two of the serums was used twice in the control experiments) in (A), or in the different types of groups based on the level of both the DEK and IL-8 in (B), (C) and (D). Asterisks show statistical comparison against the group treated only with the COVID-19 patients’ serum (D). Error bars: ±SD; n.s.: non-significant; *P < 0.05; **P < 0.01; ***P < 0.001 (Mann-Whitney test).

## DISCUSSION

NETosis is correlated with the progression and severity of COVID-19, and the search for new drugs or clinical trials of existing drugs targeting NETosis is ongoing [21, 22]. Treatment modalities such as direct inhibition of cytokines IL-1β, IL-6, and IL-8 or blocking the CXCR1/CXCR2 receptors with drugs like reparixin offer promising results [22–26]. Although the role of the secreted DEK in autoimmune disease-related NETosis was explored, to our knowledge, there is no available data on whether the same relationship applies to COVID-19.

Here, we found that in vitro NETosis induced by rDEK, at least partially, depends on the DEK-CXCR2 interaction, and reparixin reduces the DEK-induced NETosis. To our surprise, when we analyzed the COVID-19 patients’ samples, we showed that the amount of DEK in the serum of patients was lower than in healthy individuals, which was more obvious at the beginning of the disease. Unfortunately, we were not able to follow the disease outcome of the patients who participated in our study, therefore we couldn’t comment on whether the severity of the disease may also affect the level of DEK. It is intriguing that although the amount of DEK in the circulation of patients was lower than in healthy individuals, anti-DEK antibody still suppressed NETosis induced by patients’ serums. Based on our findings showing DEK mainly in the granules, but not in the nucleus or on the released DNA of neutrophils after the NETosis (Figures 3A and 3B), we propose that the presence of high levels of IL-8 as well as other cytokines may initiate the release of neutrophils’ granules that contain DEK, and subsequently secreted DEK may contribute NETosis via binding to CXCR2 receptor.

In summary, our results show DEK and IL-8 play a role in COVID-19-induced NETosis and the inhibition of CXCR1/CXCR2 receptors might be a potential therapeutic approach for the disease.

## ABBREVIATIONS

NET’s: neutrophil extracellular traps
SARS-CoV-2: severe acute respiratory syndrome coronavirus 2
COVID-19: coronavirus disease 2019
JIA: juvenile idiopathic arthritis
NE: extracellular neutrophil elastase
MPO: myeloperoxidase
NK: natural killer
hr: hour
hrs: hours
min: minute

## Acknowledgement

The authors thank Prof. Dr. Uygar Halis Tazebay for lab resources, Firdevs Meral and Leyla Dikmedaş for helping to collect blood samples, and Dilan Yoleri and Mehmet Tarık Çilingiroğlu for assisting with the experiments. This work was supported by the Scientific and Technological Research Council of Turkey (TÜBİTAK) Grant (SBAG 120S979).

## Author Contributions

AK conceived, designed, and supervised the experiments; wrote and edited the manuscript. İBK performed experiments and statistical analyses and wrote and proofread the manuscript. AY, İYC and TG performed experiments. A. Karahasan and NÇ provided human samples. TY provided resources and critically reviewed the results and manuscript. All authors have approved the published form of the manuscript.

