## Supplementary material for "NETosis Induced by Serum of Patients with COVID-19 is Reduced with Reparixin or Antibodies Against DEK and IL-8": Table S1

| **Group** | **Sample** | **DEK**  **(ng/ml)** | **IL-8**  **(ng/ml)** | **IL-6**  **(ng/ml)** | **IL-1β**  **(ng/ml)** | **MPO**  **(ng/ml)** | **Cit-H3**  **(pg/ml)** |
| --- | --- | --- | --- | --- | --- | --- | --- |
| DEK High - IL-8 High  (COVID-19) | CF 1 | 1297 | 2606.792 | 750.79 | 650.325 | 14959.04 | 17.60535 |
|  | CF 2 | 1610 | 3119.752 | 1036.57 | 1601.27 | 25840.24 | 12.25895 |
|  | CF 3 | 1596 | 3111.64 | 1134.02 | 1616.385 | 17197.76 | 4.49936 |
|  | CF 4 | 1633 | 2628.376 | 723.56 | 348.5555 | 40413.12 | 28.9707 |
|  | CF 5 | 1459 | 2816.528 | 637.819 | n.d. | n.d. | n.d. |
| DEK High - IL-8 Low  (COVID-19) | CF 6 | 1905 | 31.05672 | 0 | 75.6515 | 31709.76 | 25.2099 |
|  | AF 1 | 1393 | 110.8304 | 31.5899 | 44.4808 | 17022.64 | 0 |
|  | AF 2 | 1294 | 65.24976 | n.d. | 27.1709 | 27773.36 | 15.6637 |
|  | AF 3 | 1208 | 108.2256 | 95.9132 | 29.7685 | 34795.12 | 8.35485 |
|  | AF 4 | 1156 | 23.64288 | 4.97047 | 31.2923 | 21321.52 | 18.2433 |
| DEK Low - IL-8 High  (COVID-19) | CF 7 | 780 | 3115.168 | 846.138 | 3367.705 | 42009.28 | 38.457 |
|  | CF 8 | 581 | 3117.8 | 865.372 | 319.516 | 31416.48 | 31.44635 |
|  | CF 9 | 657 | 2797.32 | 643.04 | n.d. | n.d. | n.d. |
|  | CF 10 | 592 | 2845.192 | 444.435 | n.d. | n.d. | n.d. |
|  | CF 11 | 685 | 2749.504 | 193.546 | n.d. | n.d. | n.d. |
| Healthy (Contol) | H 1 | 1093 | 174.5056 | 5.17473 | 52.142 | 21529.12 | 12.5544 |
|  | H 2 | 1344 | 12.68496 | 0 | 52.338 | 15758.64 | 7.87825 |
|  | H 3 | n.d | 60.70472 | 0.003673 | 29.09425 | 21479.52 | 5.7961 |
|  | H 4 | 828 | 35.63936 | 0.057 | 77.219 | 23253.36 | 8.913 |

Supplementary Table 1. Level of analyzed cytokines in patient samples.

n.d: not determined.
